## Supplemental Information for "Engineered antimicrobial-derived peptides to manipulate mixed microbial systems"

^2^ current Address: Center of Excellence for Data-Driven Discovery, Department of Structural Biology, St. Jude Children's Research Hospital, Memphis, TN

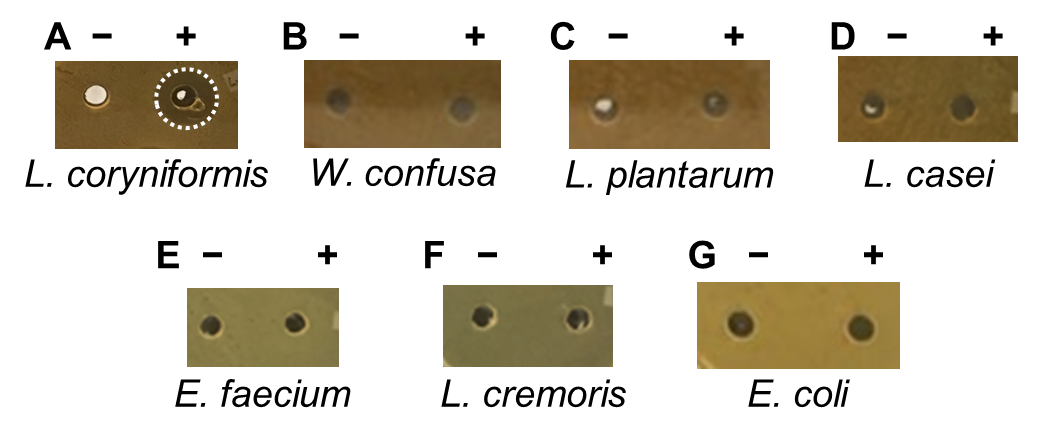

**Figure S1. Bioactivity spectrum of *P. acidilactici* UL5 spent supernatant.**

Supernatant (10 μL) from *Pediococcus acidilactici* UL-5 was used as source of pediocin PA-1 to assess bioactivity against (**A**) *Lactobacillus coryniformis* B-4390, (**B**) *Weisella confusa*, (**C**) *Lactiplantibacillus plantarum* WCSF1, (**D**) *Lacticaseibacillus* *casei* B-1922*,* (**E**) *Enterococcus faecium* NRRL B-2354*,* (**F**) *Lactococcus cremoris* MG1363, and (**G**) *E. coli* DH5α. Only *L. coryniformis* was found to be sensitive. – control; + supernatant; circle highlights the presence of zone of inhibition (ZOI).

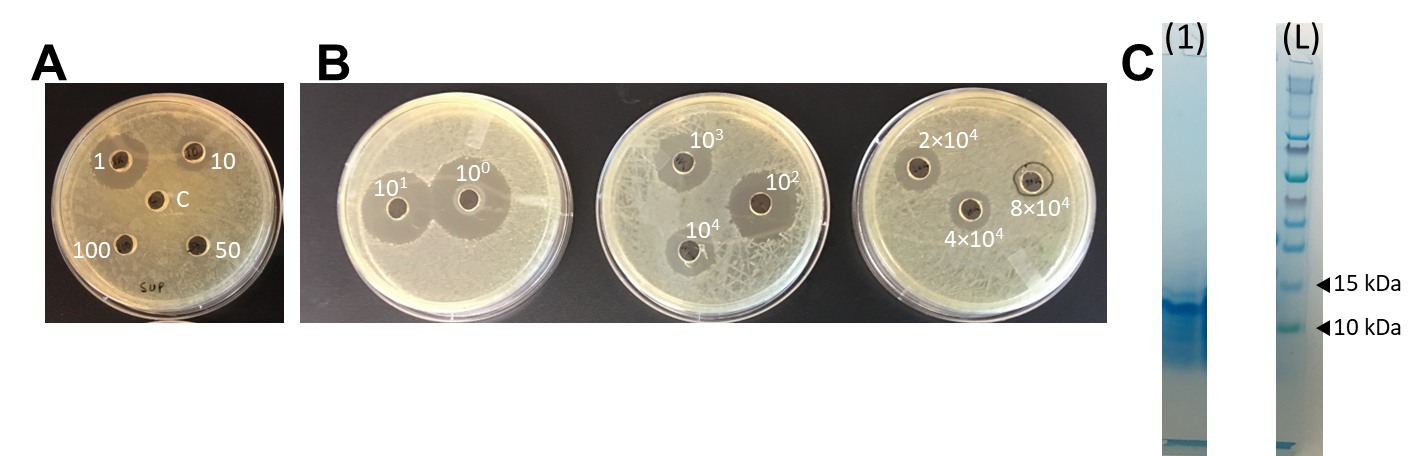

**Figure S2. Partial purification profile of pediocin PA-1 from *P. acidilactici*** **UL5.**

Bioactivity of pediocin PA-1 (10 μL) from (**A**) *P. acidilactici* UL5 culture supernatant at indicated dilutions, (**B**) partially purified pediocin PA-1 using cell adsorption-desorption method at indicated dilutions. (**C**) Reducing SDS-PAGE profile of partially purified pediocin PA-1 (lane 1) and ladder (L). C = negative control, fresh MRS.

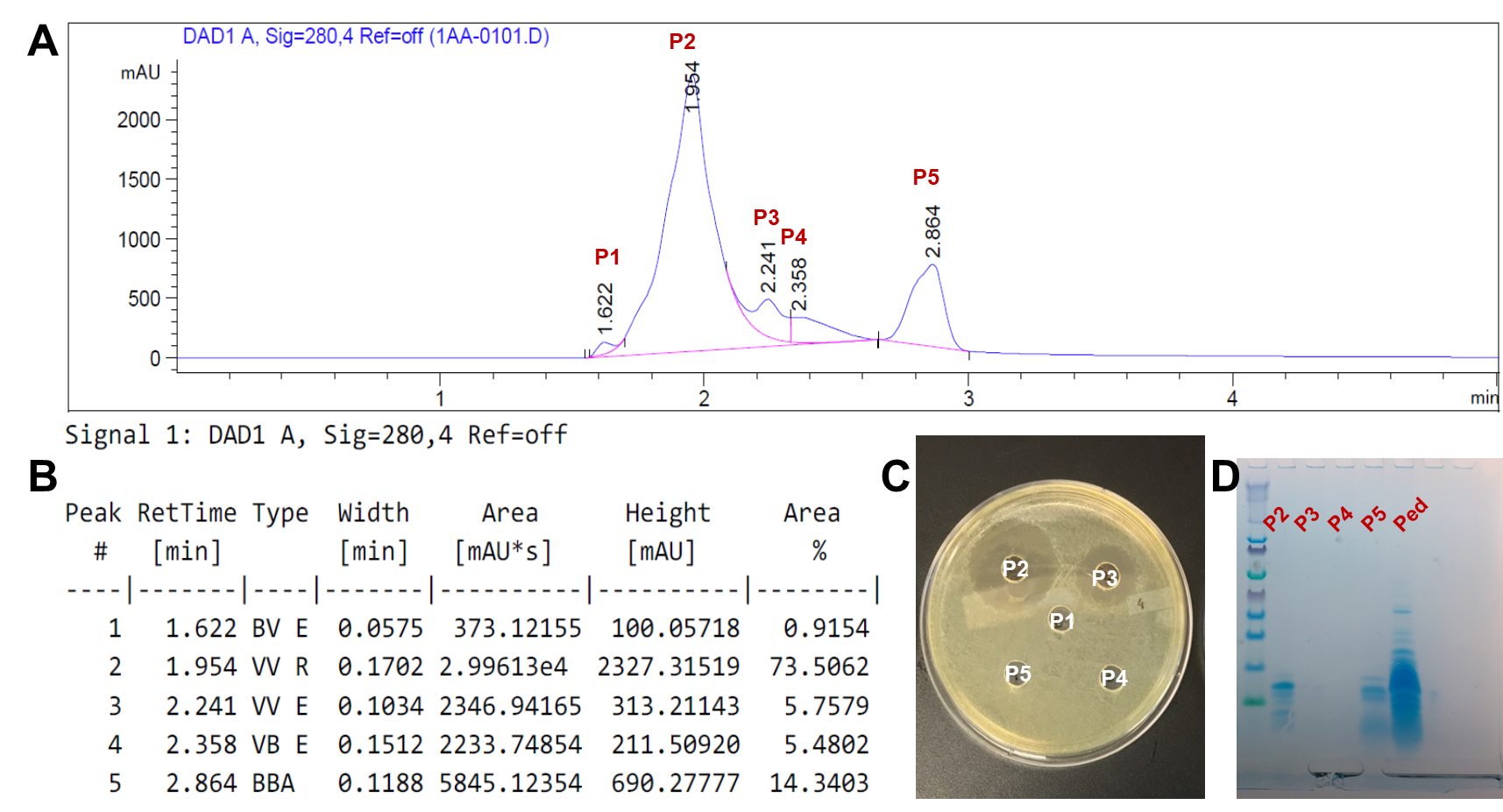

**Figure S3. Further analysis of biosynthetic pediocin PA-1.**

(**A**) Reverse phase (RP) HPLC profile of partially purified pediocin PA-1 on a C18 column. (**B**) Retention time and area of five peaks observed on the elution profile. (**C**) Bioactivity profile against *L. seeligeri.* (**D**) reducing SDS-PAGE profile of various elution fractions compared against purchased partially-purified pediocin PA-1 (MilliporeSigma).

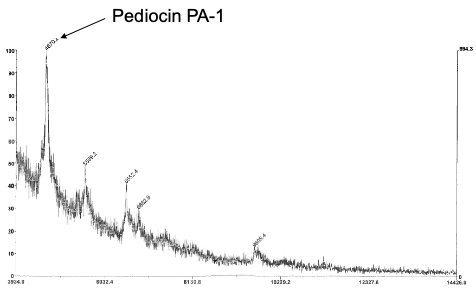

**Figure S4. MALDI-TOF profile of partially purified pediocin PA-1.**

Peptide peak with mass of 4,670.2 Da was observed in addition to peaks from other contaminating peptides in the preparation.

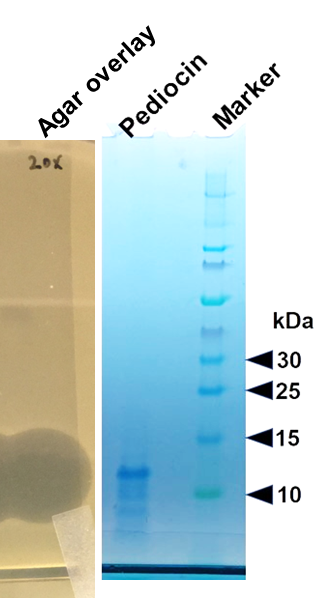

**Figure S5. Assessing activity of biosynthetic pediocin PA-1 using indicator bacterium.**

Bioactivity against *L. seeligeri* and corresponding Coomassie stained gel of pediocin PA-1 separated on non-reducing SDS-PAGE. Clearing on agar lawn suggests the ~12 kDa band corresponds to pediocin PA-1.

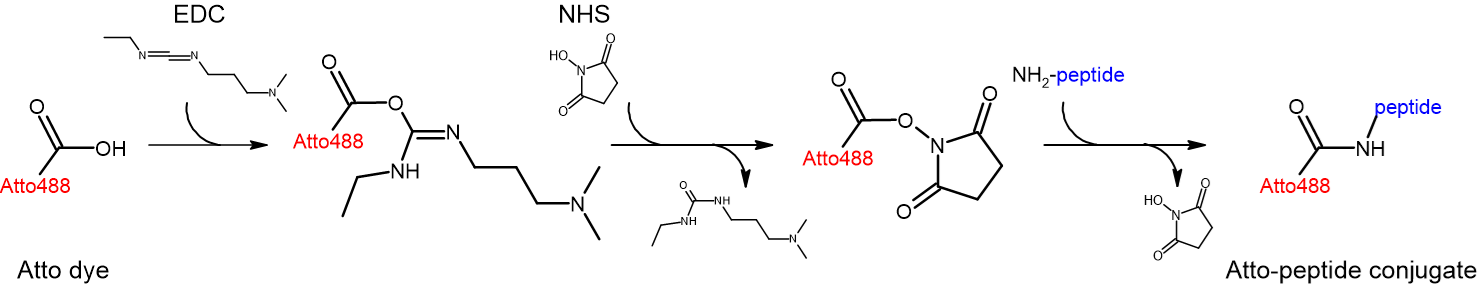

**Figure S6. Reaction scheme for EDC/NHS chemistry.**

Conjugation of pediocin PA-1 to Atto488, Atto647 dyes, or other handles were made using this approach.

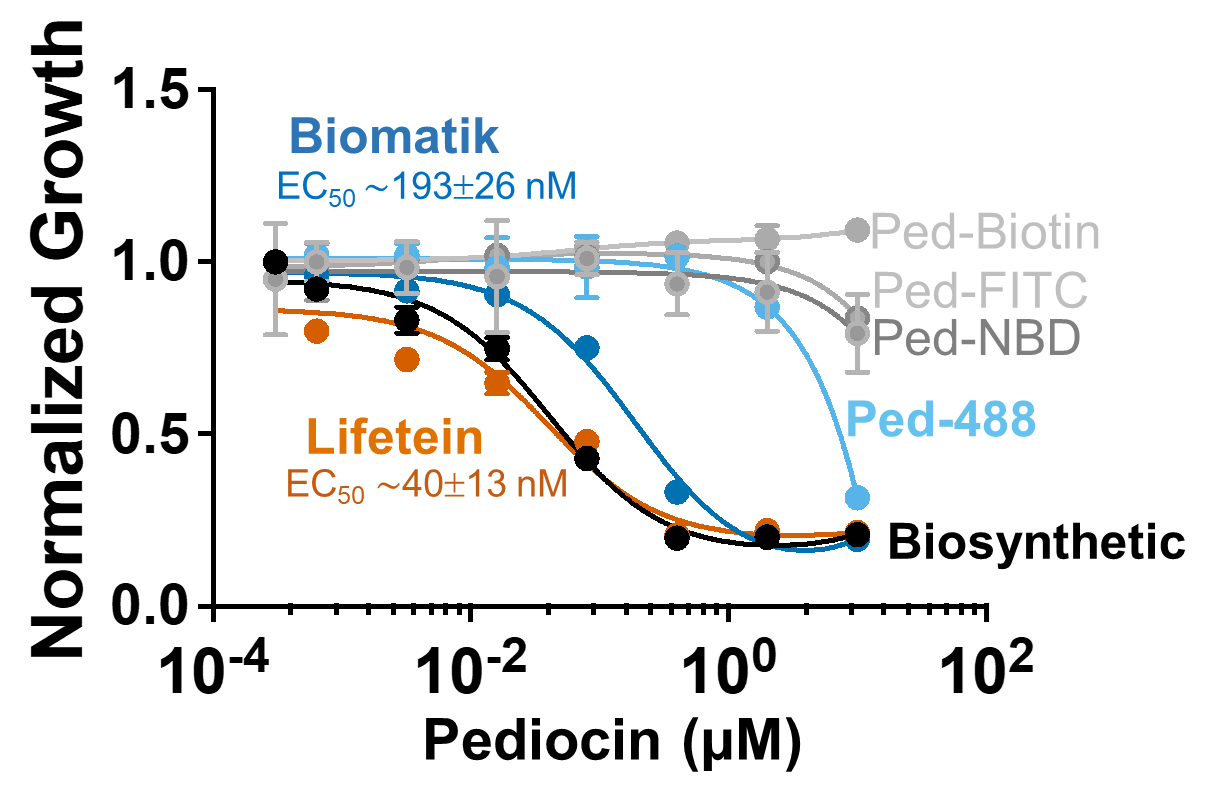

**Figure S7. Growth inhibition of *L. seeligeri* by modified and unmodified pediocin PA-1.**

The activity of Leifetein synthesized peptide (orange) was most similar to the biosynthetic preparation (black). The Biomatik preparation (dark blue) and conjugates to Atto488 (light blue), NBD, FITC, and biotin (all grey) had very low activity.

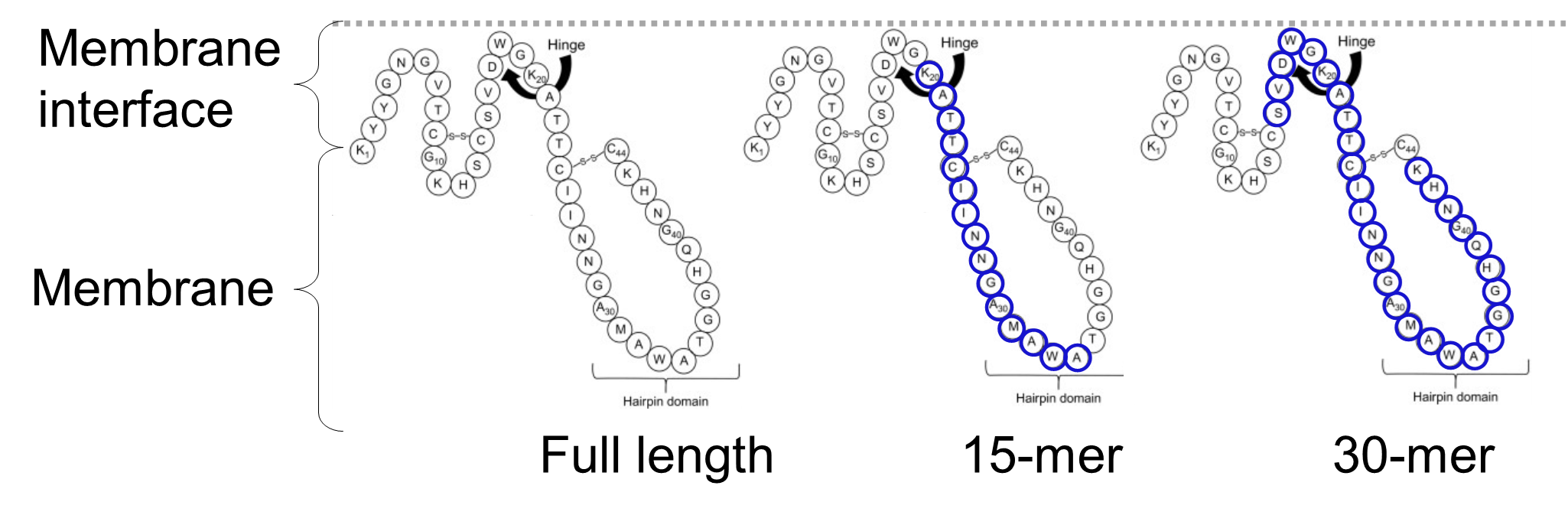

**Figure S8. Representation of pediocin PA-1, the 15-mer, and 30-mer used in this study.**

Schematic of pediocin PA-1 adapted from Espitia et al. (*Antimicrobial Food Packaging*

2016, Pages 445-454). Blue highlights residues included in truncated peptides.

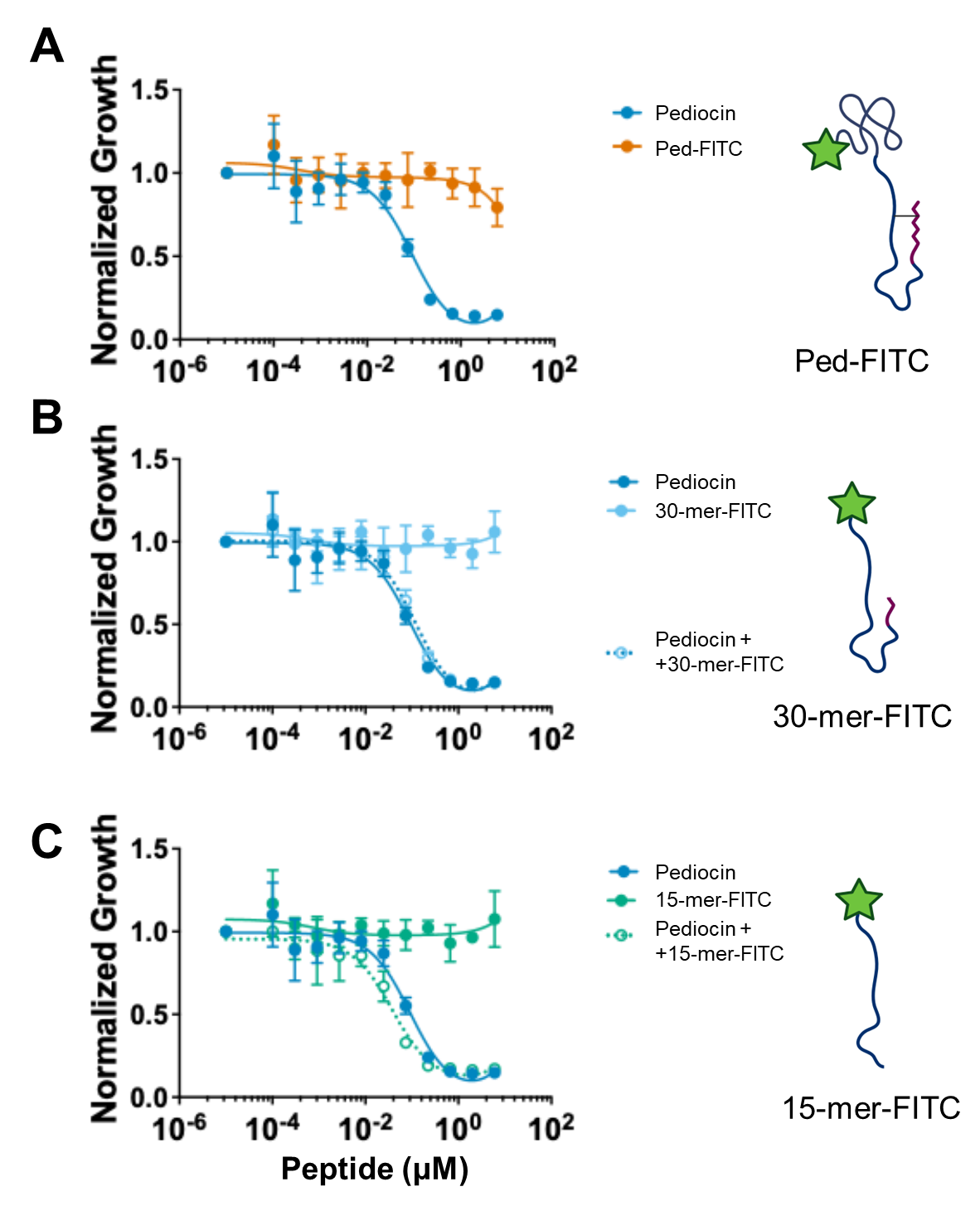

**Figure S9. Assessing bioactivity of truncated pediocins.**

Growth inhibition curve of FITC-conjugated (**A**) pediocin PA-1, (**B**) 30-mer, and (**C**) 15-mer in presence or absence of pediocin PA-1 against *L. seeligeri*.

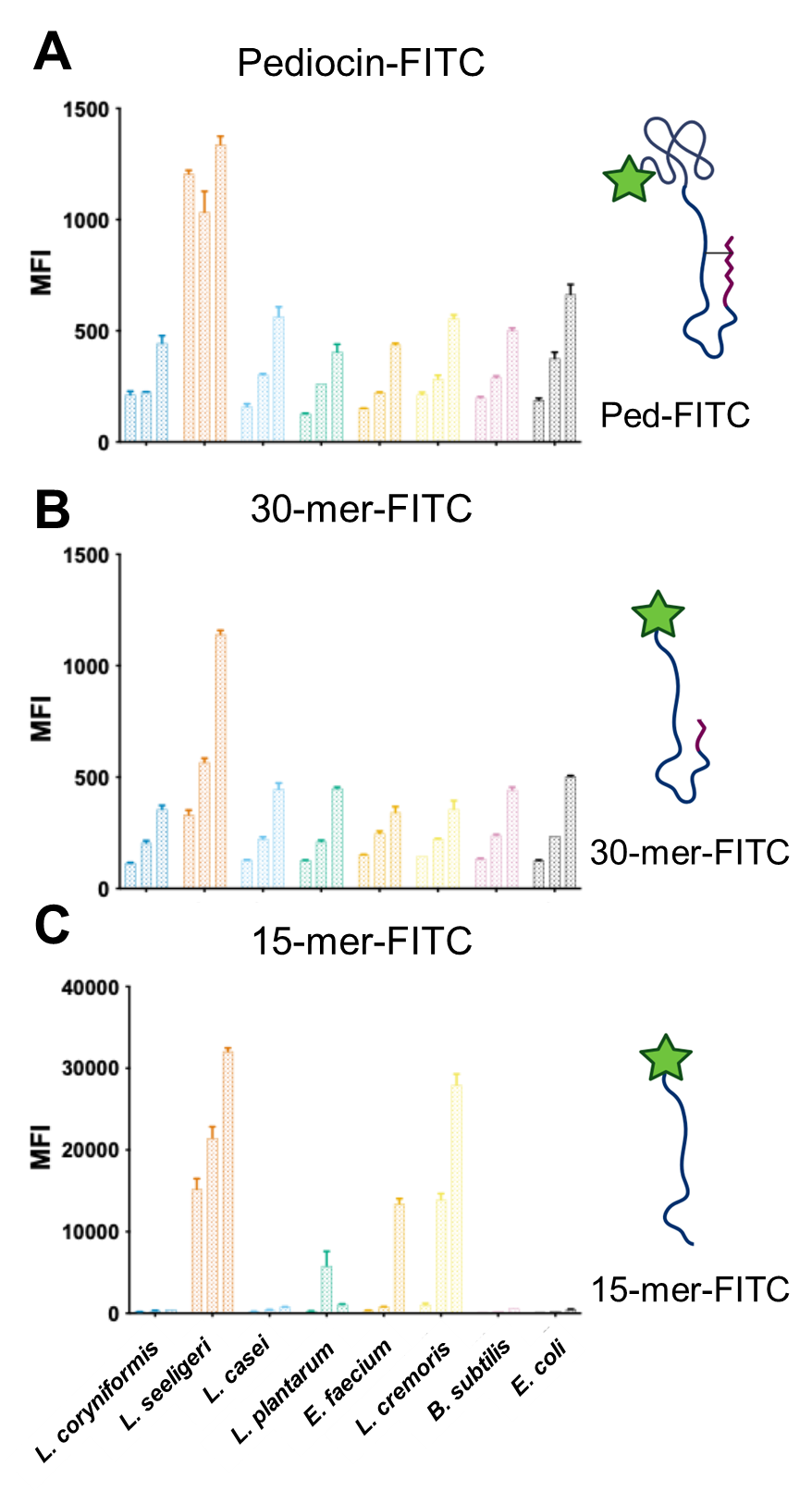

**Figure S10. Assessing pH dependence of antimicrobial activity.**

Binding spectrum of (**A**) pediocin-FITC, (**B**) 30-mer-FITC, and (**C**) 15-mer-FITC conjugates at pH 4.0 (left bars), 6.0 (middle bars), and 7.4 (right bars) to various bacteria.

**Table S1**. Apparent binding constants of pediocin PA-1 for various bacteria, where,

*MFI_max_* = maximum mean fluorescence intensity; *[S]* = peptide concentration; *n* = Hill co-efficient.

$$MFI=\frac{{MFI}_{max}{[S]}^{n}}{K_{D}^{n}+{[S]}^{n}}$$

| **Organism** | **Apparent K_D_ (nM)** | **Hill co-efficient** | **MFI_max_ (×10^3^)** |
| --- | --- | --- | --- |
| *L. coryniformis* | 46 ± 8 | 1.4 | 6.3 ± 0.3 |
| *B. subtilis* | 467 ± 167 | 0.9 | 7 ± 1 |
| *L. lactis* | >2000* | 0.7 | >2* |
| *L. casei* | >2000* | 0.7 | >2* |
| *E. coli* | >2000* | 0.9 | >2* |

* values could not be determined as the signal did not saturate.
